# A compact, portable degron tool derived from the 3′UTR of *MTCH2* for tunable degradation of proteins

**DOI:** 10.64898/2026.02.04.703731

**Authors:** Lekha E Manjunath, Saubhik Som, Ramon Boro, Anubhav Dhar, Eepsitha, Saravanan Palani, Sandeep M Eswarappa

**Author notes:** Centre for Human Genetics, Bengaluru, Karnataka, 560100, India (Lekha E Manjunath).

## Abstract

Targeted protein degradation relies on short sequence elements, termed degrons, that direct proteins to the cellular degradation machinery. Here, we report a nine-amino-acid degron (FYTVWRAFL) derived from the 3′ untranslated region (UTR) of human *MTCH2* and demonstrate its utility as a tuneable protein-degradation tool. When appended to the C-terminus of proteins, FYTVWRAFL induces rapid and robust degradation by the ubiquitin-proteasome system. Using pharmacological inhibition and genetic perturbation, we show that degradation mediated by this degron is dependent on the E3 ubiquitin ligase MDM2. Structural modeling revealed that FYTVWRAFL interacts with the hydrophobic cleft of MDM2 in a manner similar to the p53 degron. Three hydrophobic residues form the core interaction interface. By systematic mutagenesis of these residues, we generated a panel of degron variants that confer graded levels of protein stability. We demonstrate the versatility of this system by achieving tunable expression of the endogenous protein Elm1 in *Saccharomyces cerevisiae*. Collectively, our study establishes a compact, transportable, and tunable degron system as a robust toolkit for quantitative control of protein abundance across eukaryotic systems.

## INTRODUCTION

Maintenance of appropriate protein concentration, folding, localization, and activity is fundamental to cellular homeostasis (Klaips, Jayaraj et al., 2018, Labbadia & Morimoto, 2015). Among these parameters, protein abundance is tightly regulated at multiple stages of gene expression, spanning transcriptional, translational, and post-translational mechanisms. Regulated protein degradation ensures the timely removal of misfolded, damaged, and unnecessary proteins. This prevents their accumulation and the resultant cellular dysfunction.

In eukaryotic cells, the ubiquitin proteasome system (UPS) is responsible for most of the selective intracellular protein degradation (Pohl & Dikic, 2019). In this pathway, target proteins are covalently modified with ubiquitin through a series of actions by E1 ubiquitin-activating enzymes, E2 ubiquitin-conjugating enzymes, and E3 ubiquitin ligases. E3 ligases confer substrate specificity (Zheng & Shabek, 2017). Polyubiquitinated proteins are directed towards proteasomal degradation. In addition, the autophagy–lysosome system also contributes to proteostasis by clearing protein aggregates and aberrant proteins. Together, these two degradation pathways protect cells from proteotoxic stress and its pathological consequences (Pohl & Dikic, 2019).

A key determinant of selective protein degradation is the presence of short linear peptide motifs termed ‘degrons’ (Zhang, Mena et al., 2025). They are recognized by E3 ubiquitin ligases. Degrons are often located at protein termini, where they are more accessible. However, they can also reside internally and become exposed if the protein is misfolded. The modular and portable nature of degrons has been exploited experimentally as Proteolysis-Targeting Chimeras (PROTACs) (Bekes, Langley et al., 2022). In this system, a portable degron is conjugated with a molecule that specifically recognizes a protein of interest via a linker. They recruit endogenous E3 ligases to specific target proteins, inducing their degradation. PROTAC-based strategies have demonstrated encouraging therapeutic results, particularly in oncology. Multiple candidates are currently at various stages of clinical development (Bekes et al., 2022, Nalawansha & Crews, 2020). Despite these advances, there is a need for compact, portable, and genetically encoded degron modules that enable graded and context-independent control of endogenous protein levels, particularly across experimental systems and organisms.

Previously, we reported stop codon readthrough (SCR) of the mammalian *MTCH2* mRNA that encodes mitochondrial carrier homolog 2 (Manjunath, Singh et al., 2020). In SCR, translating ribosomes continue translation beyond the canonical stop codon to generate a C-terminally extended isoform, termed MTCH2xx. The proximal part of the 3′ untranslated region (UTR) encodes the C-terminal extension. We demonstrated that the SCR of *MTCH2* is important in maintaining normal mitochondrial membrane potential and normal cellular ATP levels. Importantly, MTCH2xx is markedly less stable than the canonical MTCH2 isoform, suggesting that its unique C-terminal extension encodes a degron.

In this study, we identify and characterize this C-terminal degron, demonstrate its portability and tunability, and establish that its activity is mediated by the UPS. We demonstrate the utility of this degron as a genetically encoded tool for controlled degradation of endogenous proteins. Collectively, our findings introduce a versatile degron module for precise and scalable regulation of protein abundance.

## RESULTS

### The proximal 3′ UTR of human *MTCH2* encodes a putative degron

In our previous work, we showed that MTCH2xx, one of the isoforms generated by stop codon readthrough (SCR) of *MTCH2*, is markedly less stable than the canonical MTCH2 isoform (Manjunath et al., 2020). To elucidate the mechanism underlying this instability, we examined the MTCH2xx-specific sequence (i.e., the part encoded by the proximal 3′ UTR of *MTCH2*) using the online tool Eukaryotic Linear Motif (ELM) (Kumar, Michael et al., 2024), which identifies short regulatory motifs in eukaryotic proteins. This analysis revealed a nine-amino acid stretch (FYTVWRAFL) at the C-terminus of MTCH2xx predicted to function as a degron (Fig. 1A). Notably, this sequence shares similarity with the degron in p53 that is recognized by the E3 ligase MDM2 (Brooks & Gu, 2006).

**Figure 1.**
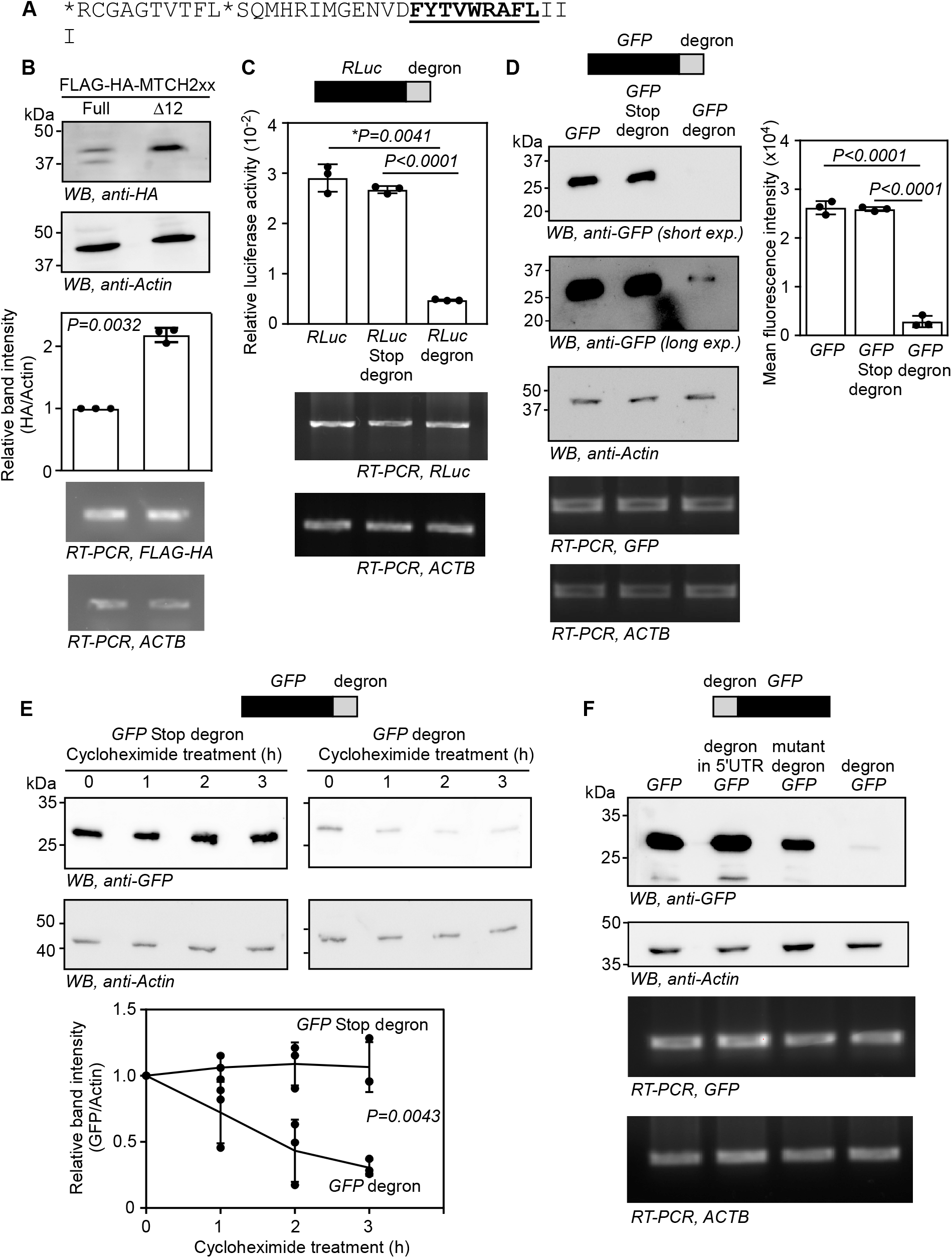
The C-terminal extension of MTCH2xx contains a transferable degron FYTVWRAFL. **(A)** Sequence of amino acids at the C-terminal extension of MTCH2xx (i.e, encoded by the proximal 3′UTR of *MTCH2* due to SCR). The putative degron sequence is highlighted. *, position of the stop codons across which readthrough happens. **(B)** Immunoblot analysis of FLAG-HA-tagged MTCH2xx expressed either in full length or without the terminal 12 amino acids, which includes the putative degron. Relative protein abundance was quantified by densitometry and is shown in the accompanying graph. **(C)** Luminescence activity of *Renilla* luciferase (RLuc) attached with the putative degron (FYTVWRAFL) at the C-terminus. Luminescence values of RLuc were normalized to those of co-transfected firefly luciferase. A control construct in which a stop codon separates the RLuc coding sequence from the degron coding sequence was included. **(D)** Immunoblot analysis of GFP expression attached with the putative degron at the C-terminus. The graph shows mean GFP fluorescence intensity measured by flow cytometry from the same cells. A control construct in which a stop codon separates the GFP coding sequence from the degron coding sequence was included. **(E)** Immunoblot analysis of GFP reporters with or without the degron following treatment with cycloheximide (100 µg ml^−1^). The graph depicts relative GFP levels quantified by densitometry. **(F)** Immunoblot analysis of GFP with the degron fused to the N-terminus. Mutant degron -(ΔF, W→A, ΔL). All experiments were performed in HEK293 cells. RT-PCR analyses in all panels show the expression of the indicated transcripts. Graphs show mean ± SD from three biological replicates. Statistical significance was determined using a ratio-paired two-tailed Student’s t-test in **(B)**, an unpaired two-tailed Student’s t-test in **(C)** and **(D)**, and two-way ANOVA in **(E)**. *, with Welch’s correction.

We expressed a FLAG-HA-tagged MTCH2xx without the C-terminal 12 amino acids (Δ12), which contains the predicted degron. Deletion of this region resulted in substantially higher protein levels than in full-length MTCH2xx in HEK293 cells. Importantly, the transcript levels of the two constructs were comparable. This observation shows that the observed difference results from post-transcriptional regulation, likely due to enhanced protein stability (Fig. 1B).

### The FYTVWRAFL degron drives the degradation of exogenously expressed proteins

We next investigated whether the nine-amino acid C-terminal motif of MTCH2xx (FYTVWRAFL) functions as a transferable degron. We appended this sequence to two reporter proteins, *Renilla* luciferase (RLuc) and green fluorescent protein (GFP). Fusion of FYTVWRAFL to the C-terminus of these reporters resulted in a marked reduction in protein abundance. This was measured by reporter activity (luminescence or fluorescence) and by immunoblotting (GFP). Insertion of a stop codon just upstream of FYTVWRAFL-coding region abolished this effect, indicating that reduced protein levels depend on the peptide sequence rather than on the corresponding RNA element. RT-PCR results revealed comparable transcript levels across all constructs, which support a post-transcriptional mechanism (Fig. 1C,D).

We performed cycloheximide-chase experiments to directly assess the effect of FYTVWRAFL on protein stability. Cells were treated with cycloheximide (100 µg ml^−1^) 20 h after transfection, and GFP protein levels were monitored over time. Levels of GFP fused to FYTVWRAFL exhibited a ∼50% reduction within 3 h of translation inhibition. However, GFP lacking the degron (GFP-stop-degron) remained stable over the same period (Fig. 1E), showing that the degron accelerated the protein turnover.

To assess the positional requirements of this motif, we fused FYTVWRAFL to the N-terminus of GFP. The degron markedly reduced GFP levels when positioned at the N-terminus. However, the degron sequence had little effect when it was placed as a part of the 5′UTR (Fig. 1F). The degron did not show appreciable effect when inserted internally (Fig. S1). Collectively, these results demonstrate that the FYTVWRAFL motif functions as a compact and portable degron capable of destabilizing exogenous proteins when positioned at either terminus of the protein.

### The ubiquitin proteosome pathway contributes to FYTVWRAFL-mediated degradation of proteins

Many degrons mediate protein degradation through the ubiquitin–proteasome system (UPS) (Zhang et al., 2025). To determine whether FYTVWRAFL belongs to this class of degrons, we inhibited the UPS using three well-characterized compounds: the proteasome inhibitors Bortezomib and MG132, and MLN7243, an inhibitor of the ubiquitin-activating enzyme E1 (Goldberg, 2012, Hyer, Milhollen et al., 2018). HEK293 cells were transfected with tagged MTCH2xx and treated with these inhibitors 20 h post-transfection. All three treatments produced a marked increase (2 to 7-fold) in MTCH2xx protein levels, indicating that the FYTVWRAFL-containing isoform is targeted for UPS-dependent degradation (Fig. 2A). In contrast, the levels of MTCH2xx lacking the degron (Δ12) did not show the same magnitude of increase upon inhibitor treatment (Fig. 2B), demonstrating that the degron is necessary for UPS-mediated degradation. Consistent with this, GFP fused to the FYTVWRAFL motif was similarly stabilized by proteasome or E1 enzyme inhibition (Fig. 2C), supporting the idea that FYTVWRAFL functions as a portable degron that uses UPS to degrade the proteins to which it is appended.

**Figure 2.**
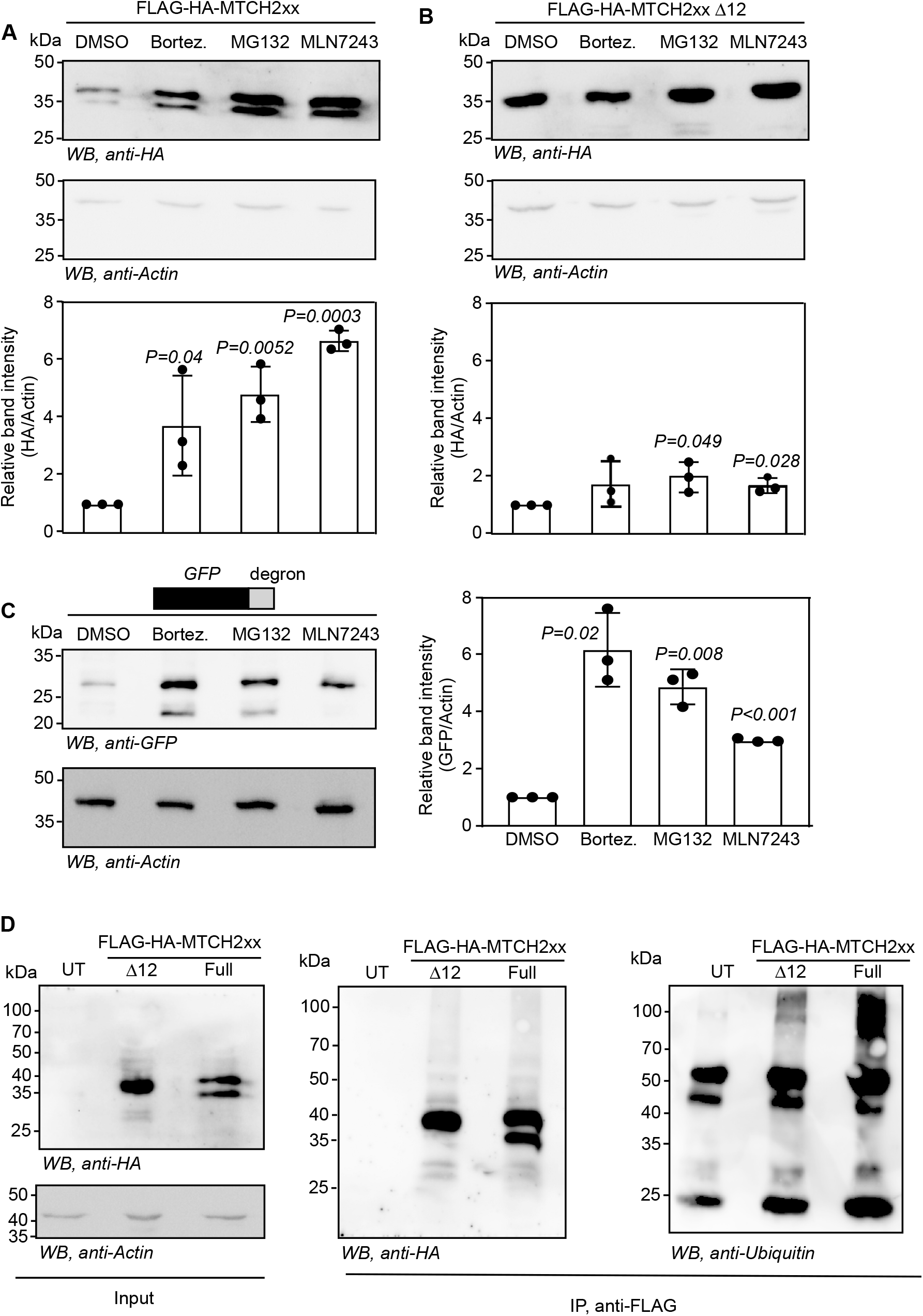
The ubiquitin–proteasome system contributes to FYTVWRAFL-mediated protein degradation. **(A–C)** Immunoblot analysis of FLAG-HA-tagged MTCH2xx expressed as the full-length protein **(A)**, without the terminal 12 amino acids **(B)**, or of GFP reporter fused to the FYTVWRAFL degron at the C-terminus **(C)**. 24 h after transfection, cells were treated for 6 h with inhibitors of the ubiquitin-proteasome pathway: bortezomib (1 µM), MG132 (10 µM), or MLN7243 (5 µM). Relative protein abundance was quantified by densitometry and is shown in the accompanying graphs. **(D)** Immunoprecipitation followed by immunoblot analysis of FLAG-HA-tagged MTCH2xx expressed with or without the C-terminal 12 amino acids. Twenty hours after transfection, cells were treated with bortezomib (1 µM) for 6 h. Equal amounts of cell lysates were subjected to immunoprecipitation using anti-FLAG beads. Immunoblotting was performed to detect FLAG-HA-tagged proteins (anti-HA) and ubiquitinated species (anti-ubiquitin). All experiments were performed in HEK293 cells. Data are presented as mean ± SD from three biological replicates. Statistical significance was assessed using a ratio-paired two-tailed Student’s *t*-test.

To further test whether degradation of MTCH2xx is ubiquitin-dependent, we expressed FLAG-HA-tagged MTCH2xx with or without the degron in HEK293 cells under conditions that yielded comparable protein levels. 20 h post-transfection, cells were treated with 1 µM Bortezomib for 6 h to inhibit the proteasome and promote accumulation of ubiquitinated intermediates. Anti-FLAG immunoprecipitation, followed by immunoblotting with anti-ubiquitin antibody, revealed multiple high-molecular-weight ubiquitinated species in both samples. Importantly, MTCH2xx, which contains the C-terminal degron, displayed substantially higher levels of ubiquitinated products compared to the construct without the degron (Fig. 2D). These results indicate that the degron enhances ubiquitin conjugation and directs MTCH2xx towards proteasome-mediated degradation.

### MDM2 mediates the degradation of FYTVWRAFL-tagged proteins

Sequence analysis using the Eukaryotic Linear Motif (ELM) resource revealed that the MTCH2xx degron (FYTVWRAFL) shares similarity with the degron present in the transactivation domain (SQET<u>FSDLWKLL</u>PEN) of p53 that is recognized by the E3 ubiquitin ligase MDM2. We examined the expression of GFP-FYTVWRAFL in MDM2 knockout HEK293 cells to test whether MDM2 contributes to the degradation of FYTVWRAFL-tagged proteins. We generated MDM2-knockout HEK293 cells using the CRISPR-Cas9 system (Fig. S2). MDM2-knockout HEK293 cells showed robust accumulation of GFP-FYTVWRAFL; however, the same fusion protein was barely detectable in parental wild-type cells (Fig. 3A). Furthermore, inhibition of MDM2 activity using a small molecule, SP141, led to a marked increase in GFP-FYTVWRAFL abundance (Wang, Qin et al., 2014)(Fig. 3B), supporting a role for MDM2 in mediating the degradation of proteins containing this motif.

**Figure 3.**
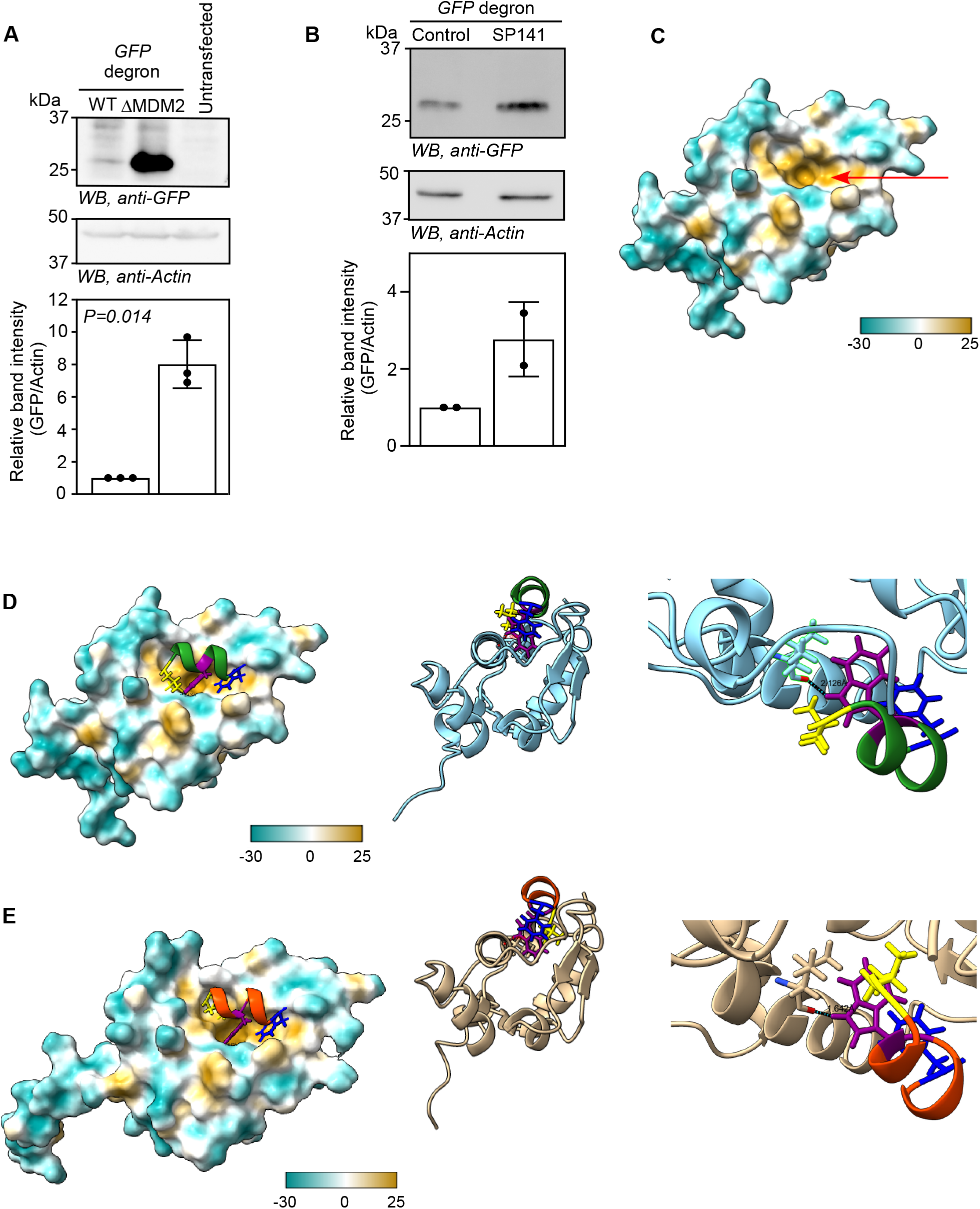
MDM2 mediates degradation of FYTVWRAFL-tagged proteins. **(A, B)** Immunoblot analysis of GFP fused to the FYTVWRAFL degron in HEK293 cells following genetic ablation of MDM2 **(A)**, or pharmacological inhibition of MDM2 using 1 μM of SP-141 for 24 h **(B)**. Data are presented as mean ± SD from two or three independent experiments. **(C)** Surface representation of hydrophobicity of the AlphaFold-predicted structure of the N-terminal domain of MDM2, highlighting the deep hydrophobic cleft (arrow) consistent with the reported crystal structure of this domain. **(D, E)** AlphaFold-predicted models of a region of MDM2 bound to the MTCH2xx degron (FYTVWRAFL) **(D)**, or the p53 degron (FSDLWKLL) **(E)**. The left panels show surface representations. Center panels show ribbon representations from another orientation, highlighting the interaction interface formed by conserved hydrophobic residues, Phe (blue), Trp (purple), and Leu (yellow), within the degron peptides. Right panels show magnified views of the binding interface. Hydrogen-bond interactions between the Trp residue of each degron and Leu60 of MDM2 are highlighted. Protein surfaces are colored according to molecular hydrophobicity, with values ranging from −30 (cyan, hydrophilic) to +25 (yellow, hydrophobic), expressed in dimensionless Molecular Lipophilicity Potential units.

To assess whether FYTVWRAFL could engage MDM2 in a manner analogous to the p53 degron, we performed structure-based modelling of the interaction. The crystal structure of the N-terminal domain of MDM2 bound to the p53 transactivation domain has been reported previously. This study revealed a deep hydrophobic cleft that accommodates the p53 degron as an amphipathic α-helix, with Phe19, Trp23, and Leu26 (NP_000537.3; <u>F</u>SDL<u>W</u>KL<u>L</u>) forming the hydrophobic interaction interface (PDB: 1YCR)(Kussie, Gorina et al., 1996). AlphaFold-based docking predicted that FYTVWRAFL binds the same hydrophobic cleft of MDM2 in an α-helical conformation. In this model, Phe1, Trp5, and Leu9 of <u>F</u>YTV<u>W</u>RAF<u>L</u> form the hydrophobic face engaging the MDM2 pocket. Furthermore, Trp of both degrons interacts with the Leu60 of the MDM2 via a hydrogen bond (2.126 Å and 1.642 Å, in case of MTCH2xx and p53 degrons, respectively). The predicted interface Template Modeling (ipTM) score for the FYTVWRAFL–MDM2 complex was 0.86, comparable to the ipTM score of 0.91 obtained for the p53 degron–MDM2 interaction using the same approach (Fig. 3C, D, and E). Together, these genetic, pharmacological, and structural analyses indicate that MDM2 recognizes the FYTVWRAFL motif and contributes to the degradation of proteins bearing this degron.

### Systematic mutations within the FYTVWRAFL degron enable graded modulation of degradation efficiency

The FYTVWRAFL motif is rich in hydrophobic amino acids (7 out of 9). The structural analysis described above identified three key hydrophobic residues within the <u>F</u>YTV<u>W</u>RAF<u>L</u> degron -phe, trp, and leu - located at positions 1, 5, and 9. These three residues form the interaction interface with the hydrophobic cleft of MDM2. To assess their significance in degron function, we generated seven variants of this degron by altering these three amino acids: ΔF (without the Phe), ΔL (without the Leu), W→A (Trp to Ala mutation), and combinations of these. These variants were appended to the C-terminus of RLuc or GFP. Protein abundance was then quantified through reporter activity assays (RLuc and GFP) and western blotting (GFP). All seven variants exhibited reduced ability to promote degradation compared to the wild-type degron, demonstrating that the identified residues are important for degron function. Importantly, the effects were graded: single-residue variants caused a modest but significant impairment of degradation, double substitutions resulted in intermediate degradation, and simultaneous alteration of all three residues nearly abolished degradation of both RLuc and GFP. These results demonstrate that the degron variants enable tunable control of the target protein.

### FYTVWRAFL degron variants enable graded control of endogenous protein levels

Strong hydrophobic patches are a hallmark of effective degrons recognized by conserved protein quality-control machinery across eukaryotes, including *Saccharomyces cerevisiae* (Hickey, Breckel et al., 2021, Johansson, Mashahreh et al., 2023). Therefore, we tested whether the FYTVWRAFL degron functions in yeast using Elm1 as a substrate. Elm1 is a septin-associated serine/threonine kinase that localizes to the mother-bud neck and regulates septin architecture and cell polarity (Bouquin, Barral et al., 2000). Loss of ELM1 results in a characteristic elongated cell morphology, which provides a sensitive phenotypic readout (Fig. 5A).

We exogenously expressed Elm1-GFP fused to the FYTVWRAFL degron or to a triple-mutant variant (described in Fig. 4). Whereas Elm1-GFP lacking the degron was robustly expressed, fusion to FYTVWRAFL rendered the protein undetectable. In contrast, the triple-mutant degron restored Elm1-GFP expression (Fig 5B). These results demonstrate that FYTVWRAFL acts as a transferable degron in yeast, similar to its function in mammalian cells.

**Figure 4.**
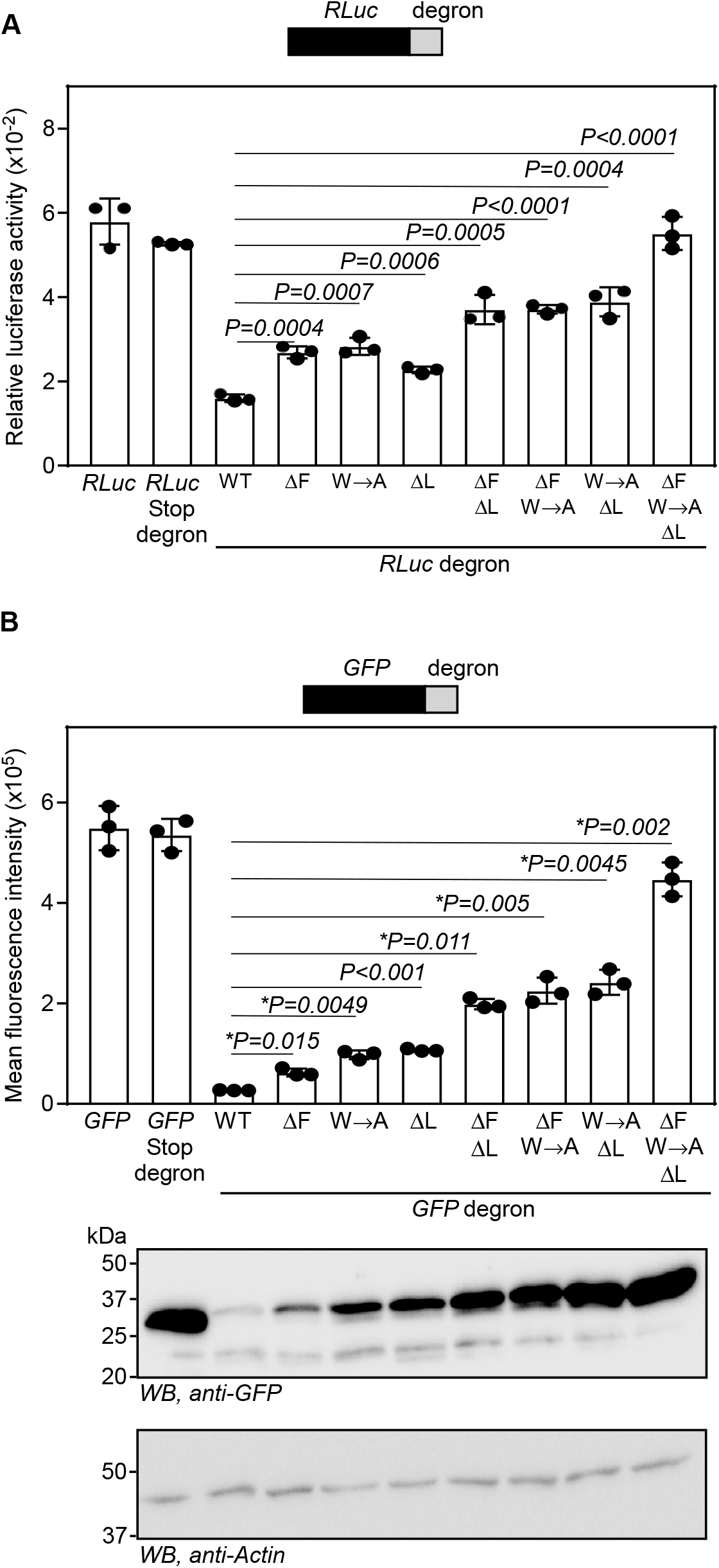
Systematic mutations within the FYTVWRAFL degron enable graded modulation of degradation efficiency. **(A)** Luminescence activity of Renilla luciferase (RLuc) fused with FYTVWRAFL degron or its variants at the C-terminus. Luminescence values of RLuc were normalized to those of co-transfected firefly luciferase. **(B)** Expression of GFP fused with the degron and its variants at the C-terminus. The graph shows mean GFP fluorescence intensity measured by flow cytometry from transfected cells. Representative immunoblots of GFP expression from the same cells are shown below. All experiments were performed in HEK293 cells 24 h post-transfection. Mutations in the degron are defined as follows: ΔF, deletion of the N-terminal phenylalanine; W→A, substitution of tryptophan with alanine; ΔL, deletion of the C-terminal leucine. Data are presented as mean ± SD from three biological replicates. Statistical significance was determined using an unpaired two-tailed Student’s t-test. *, with Welch’s correction.

**Figure 5.**
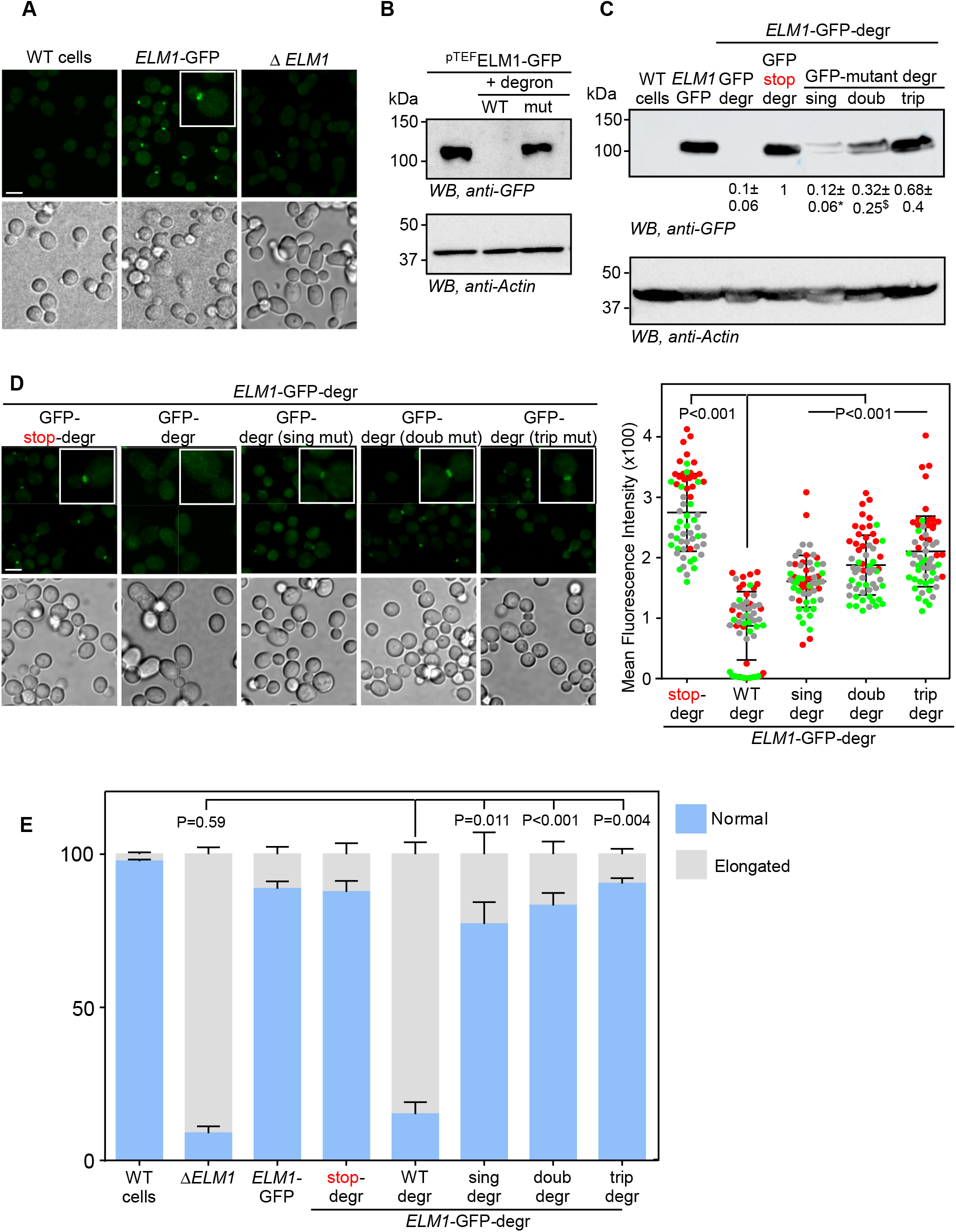
FYTVWRAFL degron variants enable graded control of endogenous Elm1 protein levels in *S. cerevisiae*. **(A)** Representative Brightfield and Fluorescence microscopy images of budding yeast cells with indicated genotypes (scale bar - 5 µm). **(B)** Western blotting image showing exogenously expressed Elm1-GFP levels. Triple mutant (ΔF, W→A, ΔL) was used as a control. **(C)** Western blotting image showing endogenously expressed ELM-1-GFP levels. Numbers below the blot indicate mean ± SD (N=3 experiments) values of density of GFP bands, normalized to the corresponding actin bands.*, *P*=0.0017 and ^$^, *P*=0.04 compared to GFP-stop-degr. **(D)** (left panel) Representative fluorescence microscopy images showing ELM1-GFP signal and localization in wildtype cells and cells expressing ELM1-GFP-degron variants (scale bar - 5 µm); (right panel) Superplot representing quantification of bud neck signal intensity of ELM1-GFP-degron variants in the indicated strains (n=20 cells per strain per replicate, N=3 replicates shown in different colors). **(E)** Stacked bar plot depicting the percentage of cells with normal or elongated morphology in the yeast strains with indicated genotypes (n>392 cells per strain per replicate, N=3). Mutants used: Single (ΔF), double (ΔF, ΔL), and triple (ΔF, W→A, ΔL). Statistical significance was determined using an unpaired two-tailed Student’s t-test.

We next examined whether FYTVWRAFL and its variants could be used to modulate endogenous protein levels. For this, we tagged GFP-FYTVWRAFL or its variants (described in Fig. 4) to the C-terminus of the endogenous *ELM1* locus. Consistent with our observations in mammalian cells, degron variants produced graded reductions in Elm1 protein abundance. The wild-type degron mediated maximum degradation, and the triple mutant exhibited substantially reduced effects. These differences were evident by Western blotting and by quantitative analysis of Elm1-GFP fluorescence at the bud neck (Fig. 5C and D).

Functionally, cells expressing Elm1-GFP-FYTVWRAFL displayed a pronounced elongated morphology, closely resembling Δelm1 cells and consistent with near-complete depletion of Elm1. In contrast, the triple-mutant degron restored both Elm1 protein levels and normal cell morphology. Notably, single- and double-mutant degrons partially restored Elm1 abundance yet were sufficient to almost fully rescue cell morphology (Fig. 5E). These findings suggest that Elm1 is present in excess at the mother bud neck and that relatively low protein levels are sufficient to sustain its cellular function, revealing a previously unappreciated tolerance of yeast cells to substantial fluctuations in Elm1 abundance.

## DISCUSSION

Eukaryotic 3′UTRs regulate gene expression, primarily via miRNA and RNA-binding protein interactions to modulate mRNA stability, localization, and translation (Mayr, 2019, West, Smith et al., 2025). An underappreciated feature of 3′UTR sequences proximal to the stop codon is their capacity to encode short C-terminal extensions when translation proceeds beyond the canonical termination site. Such extensions, generated either by termination errors or by regulated recoding of stop codons (i.e., SCR), can function as built-in safeguards that destabilize aberrant extended proteins and thereby prevent the associated proteotoxic stress (Arribere, Cenik et al., 2016). *MTCH2* exemplifies this principle; a programmed stop codon readthrough event produces an unstable readthrough isoform (MTCH2xx) that is rapidly eliminated (Manjunath et al., 2020). In this study, we identify the minimal degradation signal, i.e., the degron, responsible for this clearance and provide a compact, portable degron that can be repurposed as a tunable tool for controlling protein abundance.

Our study identifies a nine-amino-acid degron (FYTVWRAFL) that is sufficient to drive rapid degradation of fusion proteins and is functional in both mammalian cells and the budding yeast *S. cerevisiae*. In human cells, evidence from pharmacological and genetic perturbations supports a model in which MDM2, an E3 ligase, mediates degradation of FYTVWRAFL-tagged proteins. Structural modelling further predicts engagement of three key residues of the degron with the hydrophobic cleft of MDM2. This observation provided a mechanistic rationale for MDM2 dependence and a framework for rational tuning of the degron. Notably, yeast lacks MDM2 homolog, yet FYTVWRAFL remains active. This suggests that multiple E3 ligase(s) can recognize this degron. Identifying the yeast E3 ligase that recognizes this degron will be an important next step.

A key outcome of this work is a graded panel of FYTVWRAFL variants that spans a range of degradation efficiencies. This “degron series” is intended as a practical tool: it enables quantitative control of protein levels without altering transcription or translation via regulatory elements. Such control is particularly valuable when studying essential genes, where complete loss of function is lethal and partial depletion is required to uncover informative phenotypes. Using appropriate variants from the FYTVWRAFL degron series, investigators can maintain the expression levels of essential genes just enough for cells to survive while showing a phenotype that provides insights into their functions.

Beyond conditional depletion, compact degrons with defined E3 ligases are relevant to proximity-induced degradation approaches. PROTACs and related strategies utilize an E3-recruiting element to trigger ubiquitylation and degradation of target proteins (Bekes et al., 2022). Multiple degraders have advanced into advanced clinical testing. For example, vepdegestrant (an estrogen receptor degrader) in ER-positive, HER2-negative advanced breast cancer patients and BMS-986365 in metastatic castration-resistant prostate cancer patients (Campone, De Laurentiis et al., 2025)(NCT06764485).

While FYTVWRAFL is presented here primarily as a genetically encoded tag to tune protein levels, its compactness, portability, and linkage to MDM2 E3 ligase in human cells make it a potentially useful module for PROTAC development. MDM2-recruiting strategies have already been shown to support potent degradation designs (e.g., A1874), underscoring the broader translational relevance of FYTVWRAFL degron (Ficu, Niculescu-Duvaz et al., 2025, Han, Wei et al., 2022, Hines, Lartigue et al., 2019).

## MATERIALS AND METHODS

### Cell culture

HEK293 cells were cultured in Dulbecco’s Modified Eagle’s Medium (DMEM, HiMedia), supplemented with 10% fetal bovine serum (FBS, Gibco) and antibiotics (100 U/ml penicillin, 100 µg/ml streptomycin, Sigma). The cells were maintained in a humidified atmosphere at 37 °C with 5% CO2. They were authenticated by STR profiling. The cells were tested for mycoplasma contamination twice a year.

### Reagents

The following antibodies were used: anti-HA (Sigma, 11867423001), anti-Ubiquitin (Clone, UBCJ2; Enzo Life Sciences, ENZ-ABS840-0100), anti-GFP (BioLegend, 902602 or Santa Cruz, SC-9996), anti-FLAG (Clone M2; Sigma, F1804), and anti-Actin (Clone AC-15; Sigma, A3854 or Thermo Fisher Scientific, MA1-744 for yeast). Horseradish peroxidase-conjugated secondary antibodies (Invitrogen #32430; Jackson ImmunoResearch #115-035-003 and #712-035-150; Cell Signaling Technology 7076S) were used at the concentrations recommended by the manufacturers. SP-141 was from Merck (532814); Cycloheximide was from Sigma (01810-5G); Bortezomib was from Merck (5.04314.0001), MG132 was from Abcam (ab141003); MLN7243 was from Cayman Chemical (30108); Concanavalin-A was from Sigma (C2010-250MG).

### Plasmid construction

FLAG-HA-tagged MTCH2xx was cloned in the pIRESneo-FLAG/HA vector using *Not*I and *EcoR*I restriction sites, as described previously (Manjunath et al., 2020). To generate a construct lacking the putative degron, the terminal 12 amino acids of MTCH2xx were deleted (FLAG-HA-MTCH2xxΔ12). Reporter constructs expressing GFP or Renilla luciferase (RLuc) fused to the degron or its mutant variants at the C-terminus were generated in the pcDNA3.1/myc-His B vector. The coding sequence of GFP or RLuc was inserted between *Hind*III and *BamH*I, followed by the in-frame insertion of the nucleotide sequence encoding the degron or its variants between *BamH*I and *Not*I, with or without a stop codon separating them. Constructs expressing the degron at the N-terminus of GFP were also generated in pcDNA3.1/myc-His B vector by cloning the degron sequence between *Hind*III and *BamH*I, followed by insertion of GFP between *BamH*I and *Not*I.

The nucleotide sequences corresponding to the wild-type degron (FYTVWRAFL) and its mutants are given below:

Wild type: <u>TTC</u>TATACAGTG<u>TGG</u>CGCGCTTTT<u>TTA</u>

ΔF: TATACAGTGTGGCGCGCTTTTTTA

ΔL: TTCTATACAGTGTGGCGCGCTTTT

W-A: TTCTATACAGTGGCGCGCGCTTTTTTA

ΔF ΔL: TATACAGTGTGGCGCGCTTTT

ΔF W-A: TATACAGTGGCGCGCGCTTTTTTA

W-A ΔL: TTCTATACAGTGGCGCGCGCTTTT

ΔF W-A ΔL: TATACAGTGGCGCGCGCTTTT

### Luminescence- and fluorescence-based assays

HEK293 cells were seeded in 24-well plates and transfected at 75–90% confluence with 500 ng per well of plasmids encoding Renilla luciferase or GFP fused to the degron or its mutant variants. Either Lipofectamine 2000 (Invitrogen) or polyethylenimine (Polysciences) was used for transfection. For luminescence-based assays, cells were co-transfected with 100 ng per well of a firefly luciferase expression plasmid to allow normalization. Twenty-four hours post-transfection, firefly and *Renilla* luciferase activities were measured using the Dual-Luciferase Reporter Assay System (Promega) on a GloMax Explorer luminometer (Promega). For fluorescence-based assays, GFP fluorescence intensity was quantified by flow cytometry using a CytoFLEX LX instrument (Beckman Coulter) 24 h after transfection.

### Western blotting

HEK293 cells were seeded in either 24-well or 6-well plates and transfected at 75-90% confluence with 500 ng per well (24-well plates) or 2 µg per well (6-well plates) of the indicated plasmids. Transfections were performed using either Lipofectamine 2000 (Invitrogen) or polyethylenimine (Polysciences) according to the manufacturer’s instructions. Twenty-four hours post-transfection, cells were lysed in lysis buffer containing 20 mM Tris-HCl (pH 7.5), 150 mM NaCl, 1 mM EDTA, and 1% Triton X-100 supplemented with a protease inhibitor cocktail (Promega). Protein concentrations were determined using the Protein Assay Dye Reagent (Bio-Rad). Equal amounts of total protein (20-100 µg) were resolved by denaturing SDS-PAGE on 10% or 12% Tris-glycine gels and transferred onto PVDF membranes (Merck). Membranes were blocked with 5% (w/v) skimmed milk in PBS and incubated with the indicated primary antibodies, followed by horseradish peroxidase-conjugated secondary antibodies. Immunoreactive bands were visualized using Clarity ECL reagent (Bio-Rad) or Femto ECL reagent (Giri Diagnostics) and imaged using a ChemiDoc Imaging System (Bio-Rad) or Amersham ImageQuant 800 (Cytiva). Band intensities were quantified using ImageJ software.

### Cycloheximide chase assay

HEK293 cells were seeded in 6-well plates and transfected at 75-90% confluence with 2 µg per well of plasmids encoding GFP fused to the C-terminal degron, using Lipofectamine 2000 (Invitrogen). Twenty hours post-transfection, protein synthesis was inhibited by treatment with cycloheximide (100 µg/ml). Cells were harvested at the indicated time points following cycloheximide addition, and lysates were prepared for western blotting as described above.

### Immunoprecipitation

HEK293 cells were seeded in 6-well plates and transfected at 75-90% confluence with 4 µg per well of FLAG-HA-MTCH2xx or 2 µg per well of FLAG-HA-MTCH2xxΔ12 using Lipofectamine 2000 (Invitrogen). Twenty hours post-transfection, cells were treated with bortezomib (1 µM) for 6 h. Cells were then lysed in RIPA buffer supplemented with a protease inhibitor cocktail (Promega). Clarified lysates were incubated with anti-FLAG M2 affinity gel (Sigma, A2220) at 4 °C overnight with gentle tumbling. After extensive washing, immunoprecipitated proteins were eluted by boiling the beads in Laemmli sample buffer at 95 °C and subsequently analyzed by western blotting.

### RT-PCR analysis

Total RNA was isolated using RNAiso Plus (TaKaRa) according to the manufacturer’s instructions. RNA concentration was determined using a BioPhotometer (Eppendorf). Equal amounts of RNA (500 ng to 1 µg) were reverse-transcribed using oligo(dT) primers and RevertAid Reverse Transcriptase (Thermo Fisher Scientific). Gene-specific primers were then used to amplify target transcripts by PCR using Taq DNA polymerase. Primer sequences (5′ to 3′) used for amplification are listed below.

GFP: AAGTTCATCTGCACCACCG; TCCTTGAAGAAGATGGTGCG

ACTB (β-Actin): AGAGCTACGAGCTGCCTGAC; AGCACTGTGTTGGCGTACAG

FLAG-HA: GACTACAAGGACGAC; TAGCGTAATCGGGCAC

RLuc: ACTTCGAAAGTTTATGAT; TTGTTCATTTTTGAGAAC/ GGAATTATAATGCTTATCTACGTGC; CTTGCGAAAAATGAAGACCTTTTAC

### Generation of *MDM2* knockout cells

CRISPR/Cas9-mediated genome editing was employed to generate *MDM2* knockout cells. Two single-guide RNAs (sgRNAs) targeting exon 2 of the human *MDM2* gene (Ensembl gene ID: ENSG00000135679) were designed using the Benchling online tool based on the *MDM2* genomic sequence retrieved from NCBI. The first sgRNA (*MDM2*-gRNA1, 5′ CACCGCGATTGGAGGGTAGACCTG 3′) was designed to bind to the upstream region of exon 2, whereas the second sgRNA (*MDM2*-gRNA2, 5′ CACCCGAGACAAAAATACTAACCA 3′) targeted a downstream site within the same exon. Each sgRNA was cloned into the pSpCas9(BB)-2A-GFP (PX458) plasmid, which expresses *Streptococcus pyogenes* Cas9 and GFP. Two μg of each sgRNA-expressing plasmids were transfected into HEK293 cells at ∼65% confluency in a 35-mm dish using Lipofectamine 2000. At 24 h post-transfection, GFP-positive cells were isolated by fluorescence-activated cell sorting and seeded at a single-cell-per-well density in 96-well plates using a FACSAria™ II cell sorter (BD Biosciences). Populations of cells expanded from these clones were screened for the genomic deletion by PCR using primers flanking the region of expected deletion (5’ GAGTTAAGTCCTGACTTGTCTCCA 3’ and 5’ GAAGTTACGTTTGTTACGTGACTG 3’). The genomic disruption of the *MDM2* locus was confirmed by Sanger sequencing.

#### Saccharomyces cerevisiae

All yeast genetic manipulations were performed in the *Saccharomyces cerevisiae* strain ESM356 (S288C genetic background; genotype: MATa ura3-52 leu2Δ1 trp1Δ63 his3Δ200). Cells were cultured in YPD medium at 23 °C. Yeast transformations were carried out using the lithium acetate-based method as described previously (Sikorski & Hieter, 1989). Gene deletions and epitope tagging were generated by homologous recombination using PCR-derived cassettes (Fig. S3) (Janke, Magiera et al., 2004). Correct integration and strain construction were verified by colony PCR, immunoblotting, and fluorescence microscopy.

### Fluorescence Microscopy of *S. cerevisiae*

Cells were immobilized on 35-mm glass-bottom dishes (Cellvis, D35C4-20-1.5-N) coated with 6% (w/v) concanavalin A. Brightfield and GFP fluorescence images were acquired using an Olympus SpinSR spinning-disk confocal system built on a fully motorized Olympus IX83 inverted microscope. GFP excitation was achieved using a 488-nm solid-state laser, and images were captured with a Prime BSI sCMOS camera (Teledyne Photometrics). Image acquisition and analysis were performed using Fiji (ImageJ). For quantification of GFP signal intensity at the bud neck, a region of interest (ROI) was manually drawn at the bud neck, and a second ROI was drawn in a cell-free area to determine background fluorescence. Mean background intensity was subtracted from the bud-neck ROI to obtain background-corrected fluorescence values.

### Statistics

Statistical analyses were performed using either paired or unpaired two-tailed Student’s t-tests, as indicated in the figure legends. Welch’s correction was applied when variances between groups were unequal. Two-way analysis of variance (ANOVA) was used for time-course experiments assessing protein degradation. All data are presented as mean ± standard deviation (SD) from the indicated number of biological replicates. Exact N values and the statistical tests used are specified in the figure legends.

## LEGENDS TO SUPPLEMENTARY FIGURES

**Figure S1.**
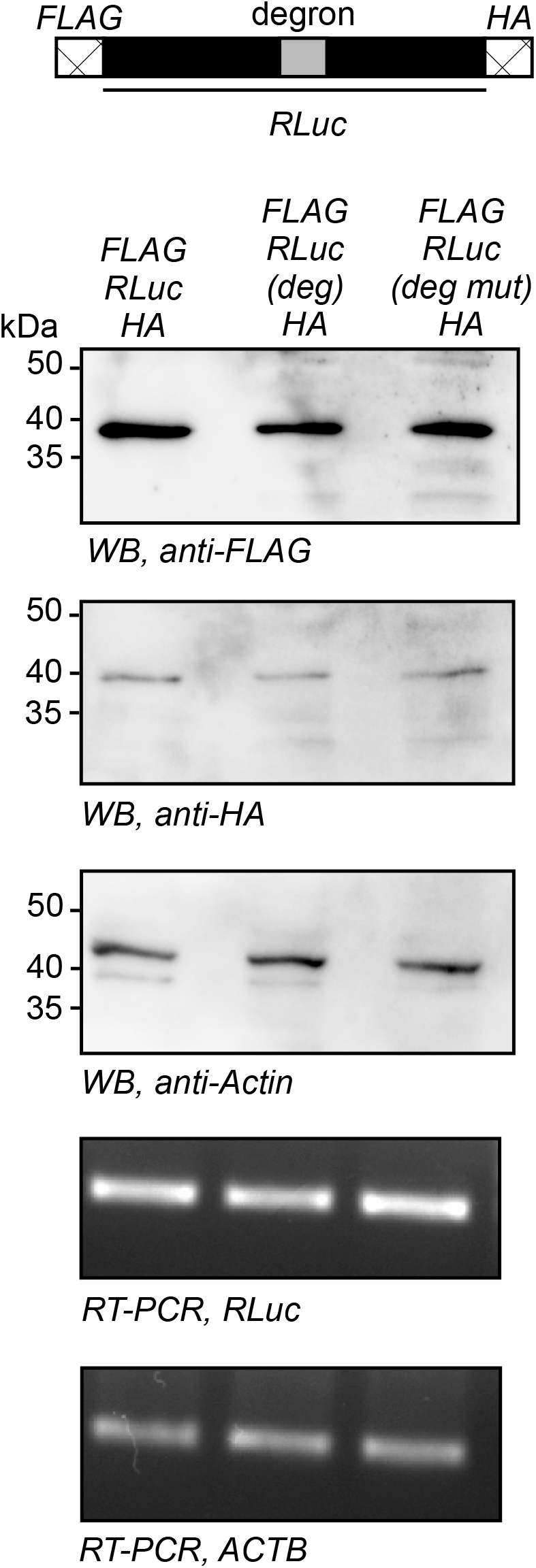
Immunoblot analysis of the expression of *Renilla* luciferase (RLuc) with internal degron. The construct was cloned in pcDNA3.1B vector. The FLAG tag was cloned between *Hind*III and *Kpn*I. RLuc was cloned between *Kpn*I and *BamH*I. The HA tag was cloned between *BamH*I and *EcoR*I. The FYTVWRAFL degron or its triple mutant (ΔF, W→A, ΔL) was inserted in the middle of the RLuc coding sequence at the *Xcm*I restriction site. These constructs were transfected in HEK293 cells. Twenty-four hours post-transfection, the samples were lysed and subjected to immunoblotting with anti-FLAG and anti-HA antibodies. The result is a representative of three independent experiments.

**Figure S2.**
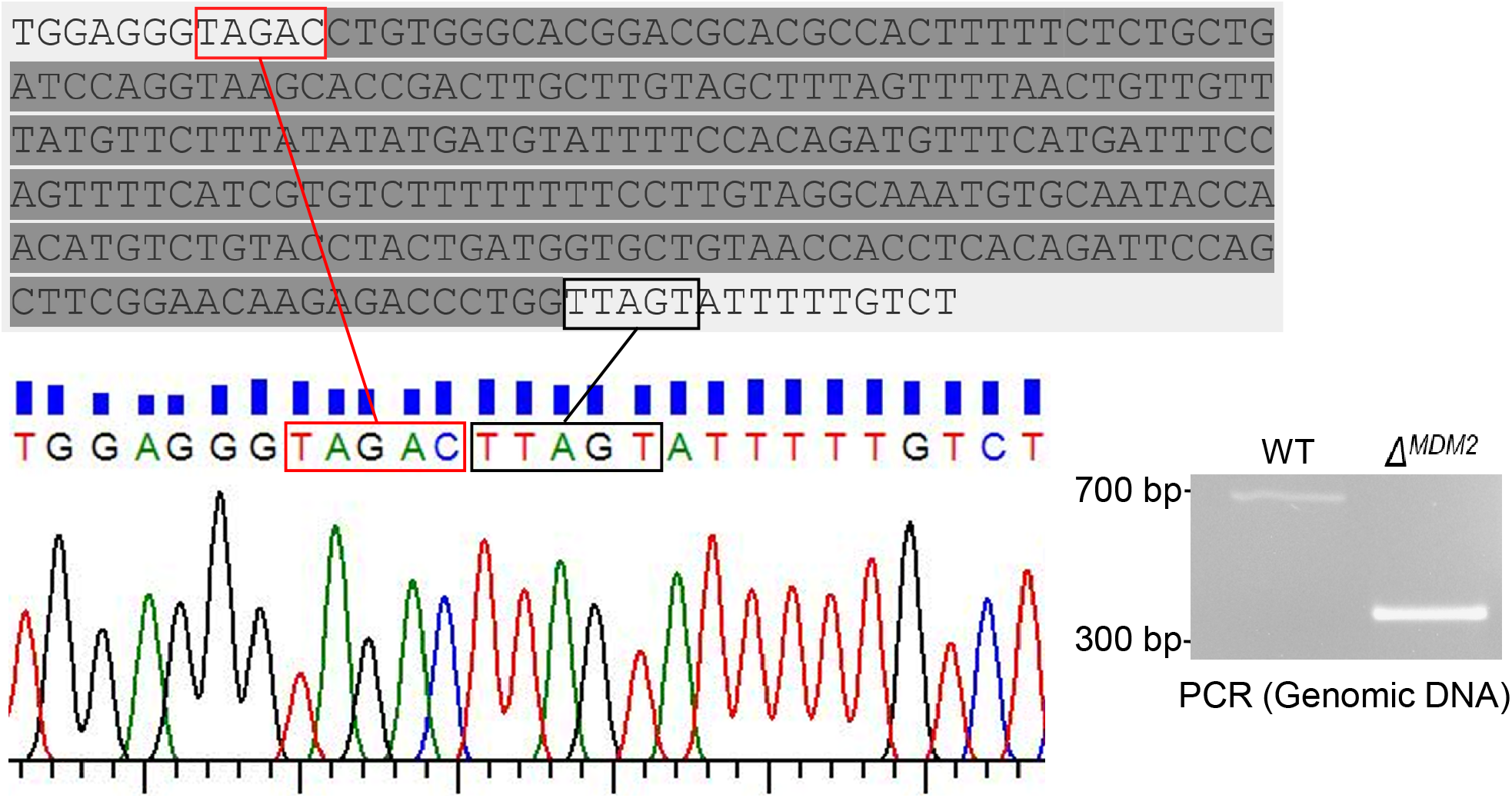
Confirmation of *MDM2* knockout HEK293 cells. Deletion was confirmed by PCR amplification from genomic DNA using primers flanking the targeted region, and by sequencing the product. Sequencing was performed using a reverse primer that generated a reverse-complement sequence. A part of this is shown as an electropherogram. The deleted part is shown in a gray background.

**Figure S3.**
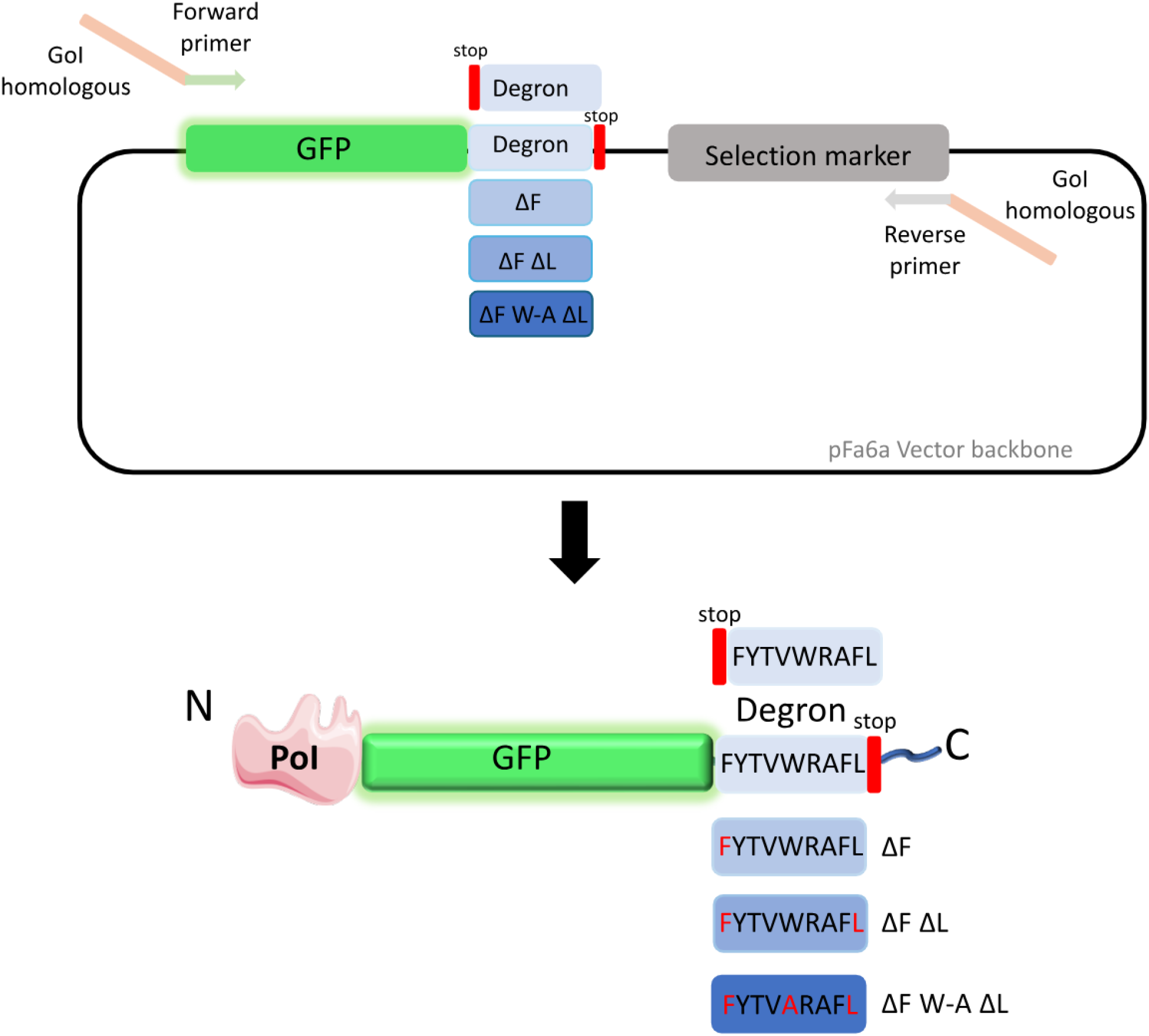
Strategy for endogenous tagging of yeast genes with GFP-degron variants. Schematic representation of the strategy used to endogenously tag yeast genes with GFP-degron variants via PCR-based homologous recombination. Briefly, tagging cassettes were PCR-amplified from the indicated plasmids using gene-specific primers containing homology arms to the target locus. The amplified cassette was transformed into competent *Saccharomyces cerevisiae* cells, where it integrated at the C-terminus of the gene of interest (GOI) through homologous recombination. Correctly modified cells were selected using the appropriate selectable marker, resulting in stable yeast strains expressing C-terminally GFP-degron-tagged proteins from their endogenous loci.

## ACKNOWLEDGEMENTS

We acknowledge the Swarnajayanti Fellowship given by the Department of Science and Technology (DST; DST/SJF/LSA-04/2019-20), grants from the Indian Council for Medical Research (EMDR/SG/12/2023-1525), Anusandhan National Research Foundation (CRG/2023/003231), and funds from EMBO Global Investigator Network, DST Funds for Improvement of S&T infrastructure, and the funds given by the Ministry of Education, India to the Indian Institute of Science. SP acknowledges funding from the Department of Biotechnology-Wellcome Trust India Alliance Intermediate fellowship (IA/I/21/1/505633). RB and AD acknowledge UGC and GATE for their fellowship.

